# Epigenetic suppression of SLFN11 in germinal center B cells in the process of the dynamic expression change during B-cell development

**DOI:** 10.1101/2020.07.30.228650

**Authors:** Fumiya Moribe, Momoko Nishikori, Hiroyuki Sasanuma, Remi Akagawa, Hiroshi Arima, Shunichi Takeda, Akifumi Takaori-Kondo, Junko Murai

## Abstract

**Background:** SLFN11 enhances cellular toxicity of genotoxic anti-cancer agents, and its important role under physiological conditions has not been appreciated yet. Somatic hypermutations and class switch recombination that can cause physiological genotoxic stress arise in germinal center B cells (GCBs). GCBs are a major origin of B-cell lymphomas that are frequently treated by cytosine arabinoside, a genotoxic anti-cancer agent.

**Objective:** To clarify the expression profile of *SLFN11* in different stages of B cells and B-cell lymphomas.

**Methods:** We analyzed the expression of *SLFN11* by mining publicly available databases for different stages of normal B cells and various types of B-cell lymphoma lines and also by performing immunohistochemical staining of human lymph nodes. We investigated the effects of two epigenetic modifiers, an EZH2 inhibitor, tazemetostat (EPZ6438) and a histone deacetylase inhibitor, panobinostat (LBH589) on *SLFN11* expression in B-cell lymphoma lines and examined the therapeutic efficacy of these epigenetic modifiers in the combination with cytosine arabinoside.

**Results:** *SLFN11* expression was specifically low in GCBs compared to non-GCBs, which was consolidated by the immunohistochemical staining for SLFN11 with human lymph nodes. *SLFN11* expression levels in B-cell lymphoma lines largely correlated to those of their normal counterparts. The epigenetic modifiers upregulated *SLFN11* expression in GCB-derived lymphomas and made the lymphomas further susceptible to cytosine arabinoside.

**Conclusions:** The expression of *SLFN11* may be epigenetically suppressed in GCBs and GCB-derived lymphomas. GCB-derived lymphomas with low *SLFN11* expression can be treated by the combination of epigenetic modifiers and cytosine arabinoside.

## INTRODUCTION

*Schlafen* (*Slfn*) family members expand in mammals. Slfn family members in mouse (*Slfn1, 2, 3, 4, 5, 8, 9, 10 and 14*) only partially overlap with those in human (*SLFN5, 11, 12, 13 and 14*) [1]. While mouse Slfns have been reported to function in immune response and lymphocyte development, functions of human SLFNs have not been well studied until recently [2-4].

SLFN11, a putative DNA/RNA helicase, has lately been analyzed from multiple aspects such as DNA damage response [5, 6], RNA cleavage [7], and defense of viral infection [8-10]. As for the role in DNA damage response, accumulative studies have shown that SLFN11 augments sensitivity to a wide range of DNA-damaging agents such as platinum-derivatives, topoisomerase inhibitors, PARP inhibitors and replication inhibitors [4-6, 11-14]. Clinical studies have proved the potency of SLFN11 acting as a predictive biomarker for the response to these drugs in lung and breast cancers [15, 16]. Mechanistically, SLFN11 binds to chromatin at stressed replication forks that are generated after DNA damage and selectively blocks fork progression, and consequently induces cell death [17, 18]. Hence, SLFN11 has a significant role to selectively eliminate cells harboring genotoxic stress.

B cells undergo gene editing mechanism at variable regions of immunoglobulin gene loci during the development and the maturation. During this process, B cells are physiologically exposed to genotoxic stress caused by somatic hypermutations and class switch recombination [19, 20]. They are introduced particularly to B cells in germinal centers (GCs) of lymph nodes by activation-induced cytidine deaminase (AID) to raise the affinity of antibodies [21]. AID further induces DNA deamination at non-targeted genes [22], thereby GCBs are presumably exposed to genotoxic stress. Hence, we hypothesized that the expression of SLFN11 needs to be well controlled during B-cell development to avoid SLFN11-dependent cell death under genotoxic stress. However, the regulatory mechanisms and functions of SLFN11 during B-cell development have not been studied almost at all, partly because SLFN11 ortholog does not exist in mice [1].

The expression level of *SLFN11* widely varies among cell types and tissues [6, 23]. The expression of *SLFN11* is largely regulated by epigenetic modifications of DNA and/or histones at its promoters and the gene body while deleterious mutations of *SLFN11* are rarely reported [24, 25]. Hence *SLFN11* expression can be activated by epigenetic modifiers such as inhibitors for DNA methyltransferases, EZH2 a histone methyltransferase, and histone deacetylases [24-26].

Based on these backgrounds, we aimed in this study to clarify the regulatory mechanisms of *SLFN11* expression and its potential roles played in B cells. We have found that *SLFN11* expression is epigenetically regulated during B-cell differentiation, and it is typically suppressed in GCBs. Moreover, epigenetic activation of *SLFN11* in lymphomas of GCB origin enhanced susceptibility to a DNA-damaging agent.

## MATERIALS AND METHODS

### Analyses of gene expression data sets

Microarray gene expression data derived from flow-sorted B-cell subsets in human bone marrow and tonsil were obtained from NCBI’s Gene Expression Omnibus database (GSE68878 and GSE69033) [27]. The exon array data were RMA normalized using R/BioC and a custom Chip Description File (CDF) [28, 29].

RNA-sequence gene expression data derived from 1001 diffuse large B-cell lymphoma (DLBCL) samples and the core set of 624 DLBCL samples were obtained from EGA (dataset id: EGA00001003600). Gene expression was measured using terms of fragments per kilobase of exon per million fragments mapped and normalized using the Cufflinks package, version 2.2.1 [30]. Quantile normalization was performed, and the data were log2 normalized.

### Immunohistochemical staining

For immunohistochemical staining analysis, we used a formalin-fixed paraffin-embedded (FFPE) lymph node sample of a resected lymph node of a cancer patient which turned out to be negative for cancer metastasis. Written informed consent was obtained from the patient for use of FFPE sample for research under the approval of the institutional ethical review board of Kyoto University Hospital. Incubation and washing were carried out at room temperature. After deparaffinization and antigen retrieval by either of Tris EDTA buffer (pH 9.0) for SLFN11 and Citrate buffer for CD20 (121°C, 5min), endogenous peroxidase activity was blocked by 0.3% H2O2 in methyl alcohol for 30 min. The glass slides were washed in PBS (6 times, 5 each min) and mounted with 1% normal serum in PBS for 30 min. Subsequently, the primary antibody (SLFN11; Santacruz: sc-515071X, mouse monoclonal IgG against amino acids 154-203 mapping within an internal region of SLFN11 of human origin, CD20; DAKO: M0755, mouse monoclonal antibody to human tonsil B cells) was applied overnight at 4 °C. They were incubated with the biotinylated second antibody diluted to 1:300 in PBS for 40 min, followed by washes in PBS (6 times, 5min). Avidin-biotin-peroxidase complex (ABC) (ABC-Elite, Vector Laboratories, Burlingame, CA) at a dilution of 1:100 in BSA was applied for 50 min. After washing in PBS (6 times, 5min), coloring reaction was carried out with DAB and Fast Red II, and the nuclei were counterstained with hematoxylin.

### Cell culture and chemical compounds

The following cell lines used in this experiment were described previously [31-33]; a germinal center B-cell-like (GCB)-DLBCL line SU-DHL6; a Burkitt lymphoma (BL) line Daudi; follicular lymphoma (FL) lines FL18, FL218, FL318. These cell lines were tested negative for mycoplasma (TaKaRa PCR Mycoplasma Detection Set; 6601), maintained in RPMI1640 (Nacalai Tesque, Kyoto, Japan) supplemented with 10 % fetal bovine serum and 1 % penicillin/streptomycin/L-glutamine and cultured at 37 °C in a humidified incubator in the presence of 5 % CO2. An EZH2 inhibitor tazemetostat (EPZ6438) was purchased from Apexbio (Boston, MA, USA). A histone deacetylase (HDAC) inhibitor panobinostat (LBH589) was purchased from Selleck Chemicals (Houston, TX, USA). The cells were seeded at 0.2-1.0 × 10^6^ cells in 2ml of medium per well in 12 well plates and treated with 5 µM tazemetostat for 4 days or 10 nM panobinostat for 16 hours before being collected for RNA extraction and western blotting. Dimethyl sulfoxide (DMSO) was used as a vehicle control.

### Reverse transcription (RT)-polymerase chain reaction (PCR)

Total RNA was extracted using RNeasy Mini kit (Qiagen, Hilden, Germany) and complementary DNA (cDNA) was synthesized using SuperScript III First-Star and Synthesis system (Life Technologies, Carlsbad, CA, USA). Quantitative RT-PCR was performed using TB Green Premix Ex Taq II (Takara). Relative gene expression was normalized to ACTB expression. The information of primers used for RT-PCR and sequencing are available (Table S1).

### Western blotting and antibodies

To prepare whole cell lysates, cells were lysed with RIPA lysis buffer system (Santacruz Biotechnology, TX, USA). Samples were mixed with tris-glycine SDS sample buffer (Nacalai Tesque) and loaded onto tris-glycine gels (BioRad). Blotted membranes were blocked with 4% bovine serum albumin (BSA) (Sigma-Aldrich. A9418) in phosphate-buffered saline (PBS) with 0.1% tween-20 (PBST). The primary antibody was diluted in 5% BSA/PBST by 1:3000 for SLFN11, and 1:10000 for Actin and acetyl-histone H3 (Lys9) (H3K9ac). The HRP-conjugated secondary antibody for mouse or rabbit (Cell Signaling, 7074S for rabbit and 7076S for mouse) was diluted in 4% BSA/PBST by 1:10000. After the membrane was soaked in ECL solution (BioRad), the blot signal was detected with luminescent image analyzer (LAS4000, GE healthcare). The mouse monoclonal anti-SLFN11 antibody (sc-515071X, 2 mg/ml, mouse monoclonal IgG against amino acids 154-203 mapping within an internal region of SLFN11 of human origin, Santacruz), the rabbit monoclonal anti-Actin antibody (12748S, rabbit monoclonal antibody to a synthetic peptide corresponding to residues near the carboxy terminus of human β-actin protein, Cell Signaling) and the rabbit monoclonal anti-acetyl-histone H3 (Lys9) (9649S, rabbit monoclonal antibody to a synthetic peptide corresponding to the amino terminus of histone H3 in which Lys9 is acetylated, Cell Signaling) were used.

### Cell viability experiments and Flow cytometry

Cell viability experiments were performed as follows: 5 × 10^4^ cells/mL viable cells were pretreated with 100 or 500 nM tazemetostat for 4 days, and 2-5 × 10^5^ cells/mL viable cells were pretreated with 5 or 10 nM panobinostat for 16 hours; 2-32 µM AraC was added to the cells for additional 24-hour incubation followed by evaluation of cell viability using Flow cytometry. Flow cytometry was performed using FACSlyrics (BD Biosciences, San Jose, CA, USA). Propidium Iodide Solution (Biolegend; 421301) was used for the evaluation of cell viability. Data were analyzed using FlowJo software (version 10.1; Tree Star Inc, San Carlos, CA, USA). Viability (%) of treated cells was defined as treated cells/untreated cells × 100. Combination index (CI) values were assessed using the CompuSyn Software (Combosyn Inc., Paramus, NJ).

### Statistical analysis

For correlation analysis, Pearson’s correlation was used and p < 0.01 was considered to be significant. For qPCR, two-sided paired t-test was used and p < 0.05 was considered to be significant.

### Data availability

All the data for the figures in this article are available from the link below. https://data.mendeley.com/datasets/kn726mznn3/

## RESULTS

### Synchronized expression of *SLFN11* with *XBP1* and the reverse expression of *SLFN11* with *PAX5*

To understand how *SLFN11* gene expression is regulated during B-cell development and maturation, we mined publicly available microarray gene expression data of various developmental stages of primary B cells derived from healthy human bone marrow and tonsil [27]. We performed correlation analyses between *SLFN11* and the other genes, and revealed that the expression of *SLFN11* was the most positively correlated with the expression of *XBP1*, a transcription factor for B-cell terminal differentiation, among transcriptional regulators, while it was the most inversely correlated with *PAX5*, a master regulator of B-cell development (Fig 1A left, Table S2) [34]. Stage-wise plotting of the data revealed that *SLFN11* expression was almost reverse of *PAX5* expression through the developmental stages (Fig 1A right). The data arranged from premature to differentiated B-cells showed that the sequential pattern of *SLFN11* expression was almost parallel to the expression of *PRDM1* and *XBP1*, both of which are key transcription factors for B-cell terminal differentiation (Fig 1B) [21, 35]. On the other hand, other *SLFN* family members (*SLFN5, 12, 13* and *14*) did not show comparable correlation with *PAX5, PRDM1 or XBP1* (Fig S1A-B). Thus, *SLFN11* but no other *SLFNs* expression can be controlled during B-cell development under the same regulatory system for *PRDM1* and *XBP1*.

**Fig 1.**
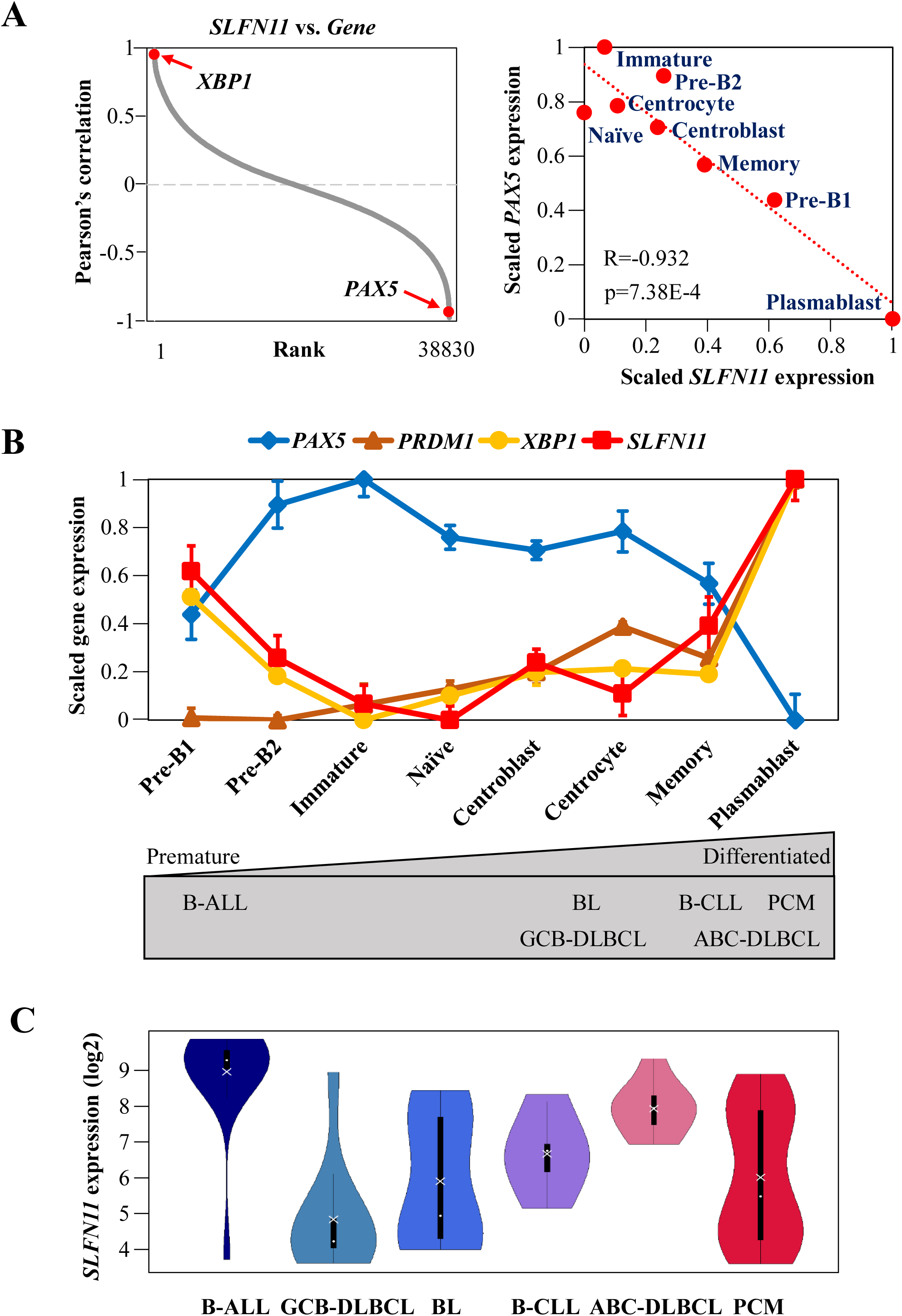
*SLFN11* expression is differentially regulated during B-cell development. (A) Left: Pearson’s correlation between *SLFN11* and all the other genes. The genes are ordered from the highest correlation (left) to the lowest correlation (right). Right: microarray gene expression plot of *SLFN11* and *PAX5*. Precursor (Pre)-B1 cells, precursor (Pre)-B2 and immature B cells are were taken from human bone marrow (n=5), and naïve B cells, centroblasts, centrocytes, memory B cells and plasmablasts were taken from human tonsil (n=6). Pearson’s correlation (R), P-value (p) and approximation straight line (a red dotted line) are shown. (B) Microarray gene expression profile (log2) of selected genes (*PAX5, PRDM1, XBP1, SLFN11*) in human B cells from bone marrow and tonsil. Dots correspond to group means ± SE (standard error). The gray figure below represents the developmental stages of the origins of B-cell lymphomas. (C) mRNA expression (log2) of *SLFN11* in B-cell derived cancer cell lines (B-ALL, GCB-DLBCL, BL, B-CLL, ABC-DLBCL, PCM). Each dot and cross correspond to the group median and mean, respectively. B-ALL, B-cell acute lymphoblastic lymphoma; GCB-DLBCL, germinal center B-cell like-diffuse large B cell lymphoma; BL, Burkitt lymphoma; B-CLL, B-cell chronic lymphocytic leukemia; ABC-DLBCL, activated B-cell like-diffuse large B cell lymphoma; PCM, plasma cell myeloma.

Next, we analyzed the public database of Cancer Cell Line Encyclopedia (CCLE) [5] to examine *SLFN11* expression levels across 79 B-cell lymphoma lines derived from different histologic subtypes: B-ALL (B-cell acute lymphoblastic lymphoma), GCB-DLBCL (germinal center B-cell like-diffuse large B cell lymphoma), BL (Burkitt lymphoma), B-CLL (B-cell chronic lymphocytic leukemia), ABC-DLBCL (activated B-cell like-diffuse large B cell lymphoma) and PCM (plasma cell myeloma). According to the origins of B cells, subtypes can be arranged from premature to differentiated B-cell lymphoma and are corresponded to their normal counterparts as shown (Fig1B below) as the origins of B-ALL, GCB-DLBCL, B-CLL and ABC-DLBCL are regarded as pre-B, centroblast / centrocyte, memory and plasmablast, respectively. GCB-DLBCL cell lines had the lowest expression levels of *SLFN11* while B-ALL and ABC-DLBCL expressed higher levels of *SLFN11* than the others (Fig 1C). Thus, *SLFN11* expression levels in these cell lines were considered to reflect those in their normal counterparts.

### Suppression of SLFN11 expression in germinal center B cells

To validate the findings of the dynamic change of SLFN11 expression during B-cell differentiation, we examined the protein expression of SLFN11 in human normal lymph node tissue. We carried out immunohistochemical staining (IHC) using an anti-CD20 antibody (a marker of B cells) and an anti-SLFN11 antibody (Fig 2). SLFN11 expression was depleted in most of the B cells in GCs whereas cortex regions, which contain non-GCBs and other cell types such as T cells and macrophages, were enriched with SLFN11 expression. These results are consistent with our findings that B cells in GCs (normal centroblasts and centrocytes, and GCB-DLBCL cells) are low in SLFN11 expression. Collectively, SLFN11 expression is tightly regulated along with B-cell development and notably, is suppressed in GCBs.

**Fig 2.**
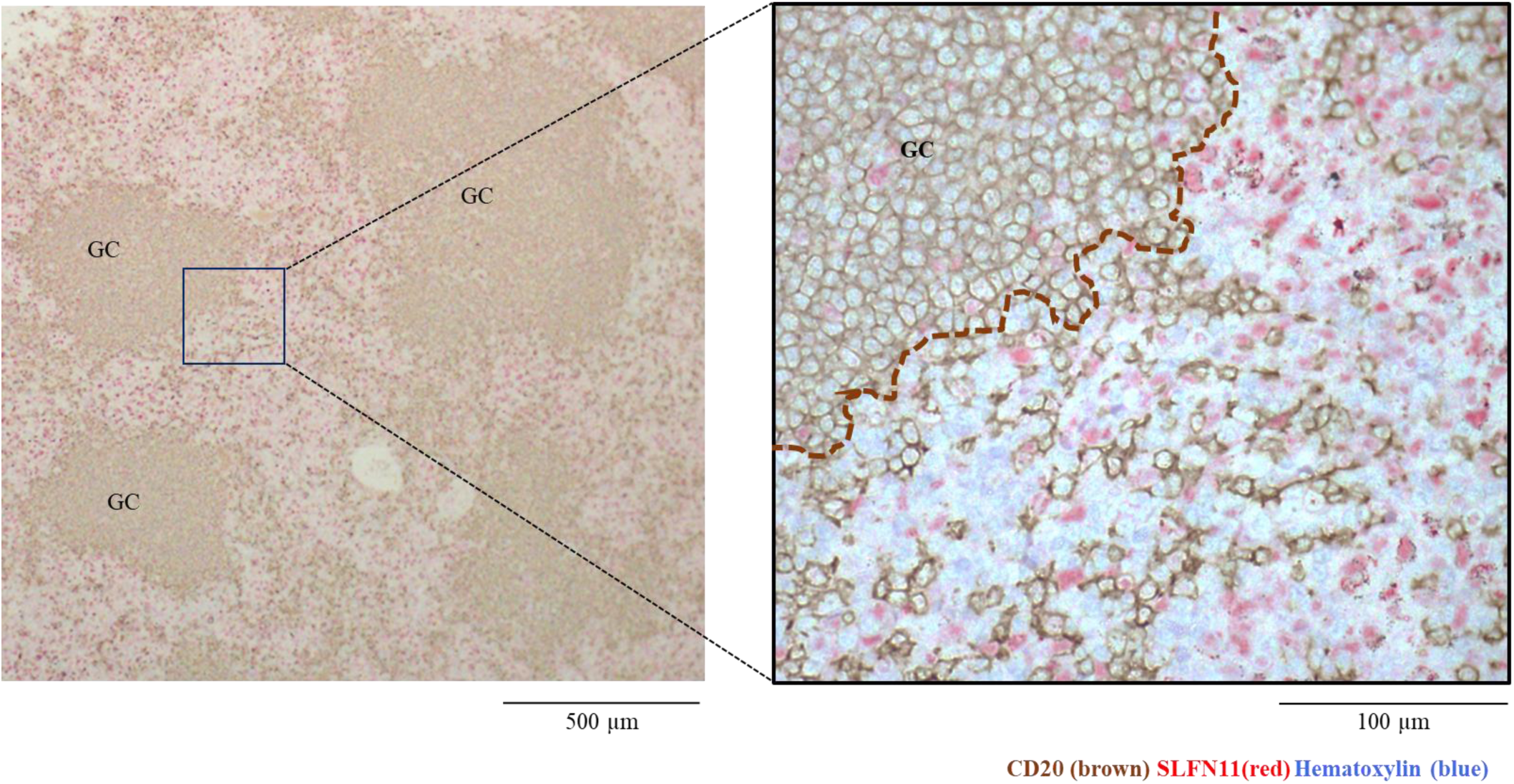
SLFN11 protein is depleted in germinal centers. Immunohistochemical staining of human lymph node tissue. SLFN11 was stained with Fast RedII (red) and CD20 was with DAB (brown). CD20 enriched regions are considered to be germinal centers. GC: germinal center

### The expression of *SLFN11* follows the expression of marker genes for ABC-DLBCL

To further study the regulatory system of *SLFN11* expression during B-cell development, we focused on the distinct transcriptional characterization between GCB-DLBCL and ABC-DLBCL. In the clinic, the high expression of BCL6, a transcriptional repressor required for GC formation, is used as an indicator of clinical diagnosis for GCB-DLBCL [36]. Consistently, GCB-DLBCL cell lines in CCLE database exhibited high *BCL6* expression whereas ABC-DLBCL cell lines had low *BCL6* expression (Fig 3A). The expression of *SLFN11* has a significantly negative correlation with that of *BCL6* across the DLBCL cell lines, suggesting that the expression of *SLFN11* can positively and reversely follow the marker genes of ABC-DLBCL and GCB-DLBCL, respectively.

**Fig 3.**
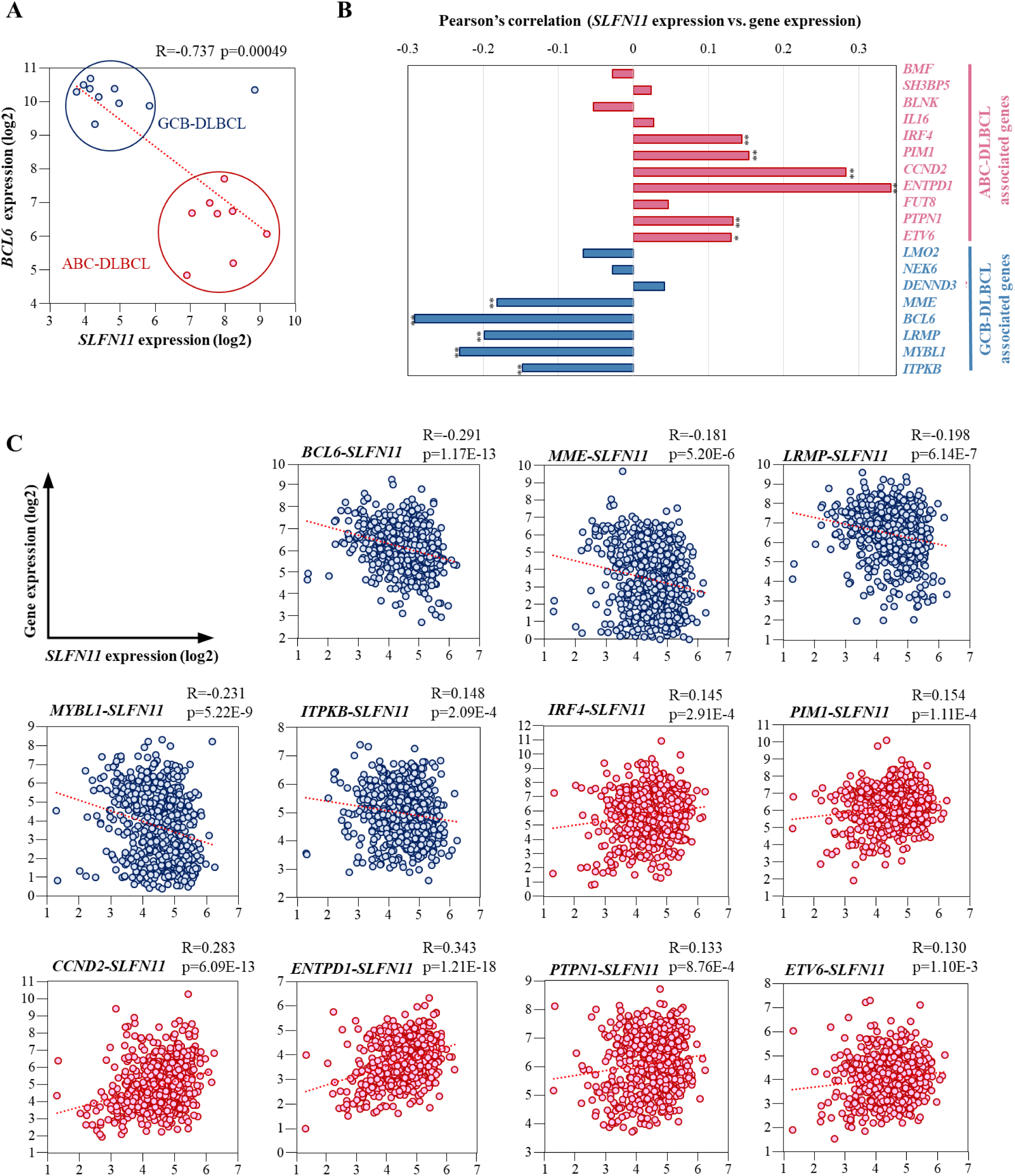
Correlation of the expression of *SLFN11* and ABC- and GCB-DLBCL associated genes in human DLBCL. (A) mRNA expression plot of BCL6 and *SLFN11*. X-axis represents *SLFN11* mRNA expression (log2). Y-axis represents *BCL6* mRNA expression (log2). The red dots are ABC-DLBCL derived cell lines and the blue dots are GCB-DLBCL derived cell lines. Pearson’s correlation (R), P-value (p) and approximation straight line (a red dotted line) are shown. (B) Pearson’s correlation of *SLFN11* expression level with DLBCL subtypes genes. ABC-DLBCL-associated genes consist of 11 genes (*BMF, SH3BP5, BLNK, IL16, IRF4, PIM1, CCND2, ENTPD1, FUT8, PTPN1, ETV6*). GCB-DLBCL-associated genes consist of 8 genes (*LMO2, NEK6, DENND3, MME, BCL6, LRMP, MYBL1, ITPKB*). *p < 0.01, **p < 0.001 (C) mRNA expression plot of DLBCL subtype genes and *SLFN11*. X-axis represents *SLFN11* mRNA expression (log2). Y-axis represents DLBCL subtype associated mRNA expression (log2). Pearson’s correlation (R), P-value (p) and approximation straight line (a red dotted line) are shown.

We then investigated the expression of *SLFN11* in DLBCL tissues from patients using the database established by Reddy et al. [37]. In addition to the *BCL6*, a set of genes has been reported to be distinctively expressed in these DLBCL subgroups and useful to classify DLBCL into either subgroup [37, 38]. The set of genes includes eleven ABC-DLBCL associated genes and eight GCB-DLBCL associated genes (Fig 3B). Correlation analyses of mRNA expression between *SLFN11* and these DLBCL-associated genes revealed that *SLFN11* expression was overall positively and negatively correlated to the expression of ABC-DLBCL- and GCB-DLBCL-associated genes, respectively. Eleven out of the listed 18 genes had significant correlations (Fig 3B-C). These results indicate that *SLFN11* expression is controlled under the gene expression program that can characterize GCB-DLBCL from ABC-DLBCL.

### *SLFN11* expression is epigenetically repressed in GCB-DLBCL

As epigenetic modifications are profoundly involved in the GCB-specific gene regulation [39-41], we hypothesized that the expression of *SLFN11* gene is also epigenetically downregulated in GCBs, like many other genes that are specifically repressed in GCBs. To explore the regulatory mechanisms of *SLFN11* gene expression in GCBs, we selected two epigenetic modifiers, an EZH2 inhibitor tazemetostat (EPZ6438) and a histone deacetylase (HDAC) inhibitor panobinostat (LBH589), both of which have been reported to upregulate the expression of genes that are specifically suppressed in GCBs [42, 43].

We first evaluated the effect of these epigenetic modifiers on the expression levels of selected GCB- and ABC-DLBCL-associated genes across the six GCB origin lymphomas: GCB-DLBCL cell line SU-DHL6, BL cell line Daudi, and FL cell lines FL18, FL218 and FL318. By quantitative RT-PCR, we found that both of these epigenetic modifiers largely upregulated ABC-DLBCL-associated genes and rather downregulated GCB-DLBCL-associated genes in the GCB origin lymphomas (Fig 4A). Under the same conditions, *SLFN11* gene expression was upregulated by both epigenetic modifiers (tazemetostat and panobinostat) across all the cell lines examined (Fig 4A). We confirmed the elevated SLFN11 expression at protein level in FL218 and FL318 cells after the treatment of either epigenetic modifier, which was consistent with the expression elevation measured by qPCR (Fig 4B). We failed to detect SLFN11 protein in the other cell lines probably because the basal expression levels of SLFN11 were too low to be detected in these cells. Based on these results, we conclude that SLFN11 is suppressed by histone methylation whereby ABC-DLBCL-associated genes are also suppressed in GCB origin lymphomas.

**Fig 4.**
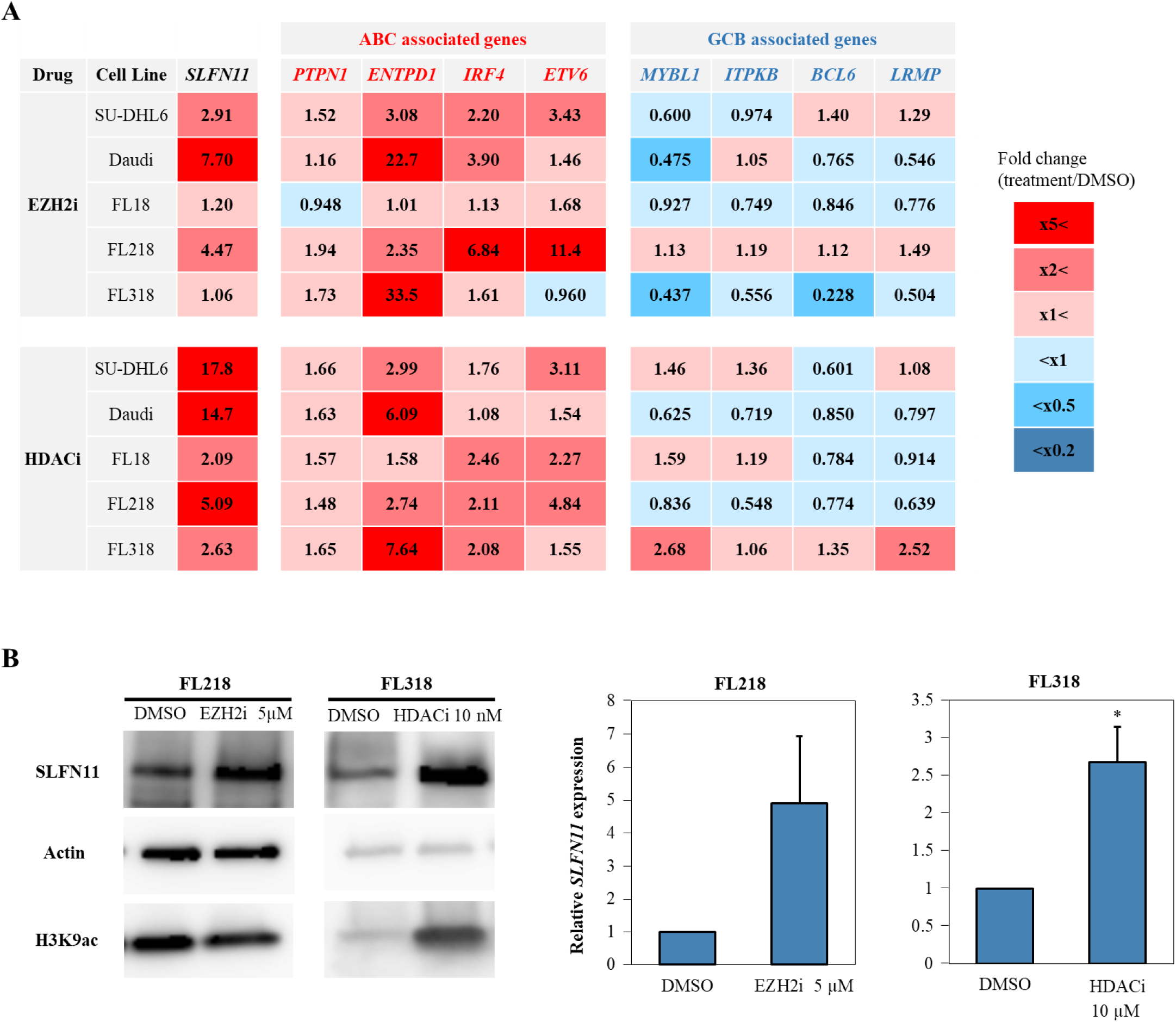
Tazemetostat/panobinostat upregulate the expressions of *SLFN11* and ABC-DLBCL associated genes in GCB lymphomas. Cells were treated with tazemetostat (5 µM, 4 days) or panobinostat (10 nM, 16 hours) and the gene expression levels were measured by quantitative RT-PCR. (A) Heatmap of fold changes (treated/untreated) of ABC- and GCB-DLBCL-associated genes. Results are the average of three independent experiments. (B) Left: western blotting of SLFN11 in FL218 treated with tazemetostat and FL318 treated with panobinostat. Right: bar graphs of *SLFN11* mRNA expression levels normalized to untreated samples. Results are the average of three independent experiments with ± SD. *p < 0.05 (two-sided paired t-test).

### Epigenetic upregulation of *SLFN11* renders GCBs more susceptible to cytosine arabinoside

To determine whether epigenetic upregulation of *SLFN11* expression enhances the sensitivity of GCB origin lymphomas to cytotoxic chemotherapy, we examined the synergistic effect of cytosine arabinoside (AraC), a replication inhibitor commonly used in lymphomas, on SU-DHL6 cells with or without pretreatment of EZH2 inhibitor (tazemetostat) or HDAC inhibitor (panobinostat). We found that the pretreatment with these epigenetic modifiers enhanced cell susceptibility to AraC in SU-DHL6 cells (Fig 5A-B left). Combination index (CI) values, a commonly used parameter to evaluate the synergistic effect, revealed that the concurrent use of AraC with these epigenetic modifiers has synergistic effects at various doses (Fig 5A-B right). These results indicate that the combination of the epigenetic modifiers with AraC can improve the drug response in GCB origin lymphomas in part by enhancing SLFN11 expression.

**Fig 5.**
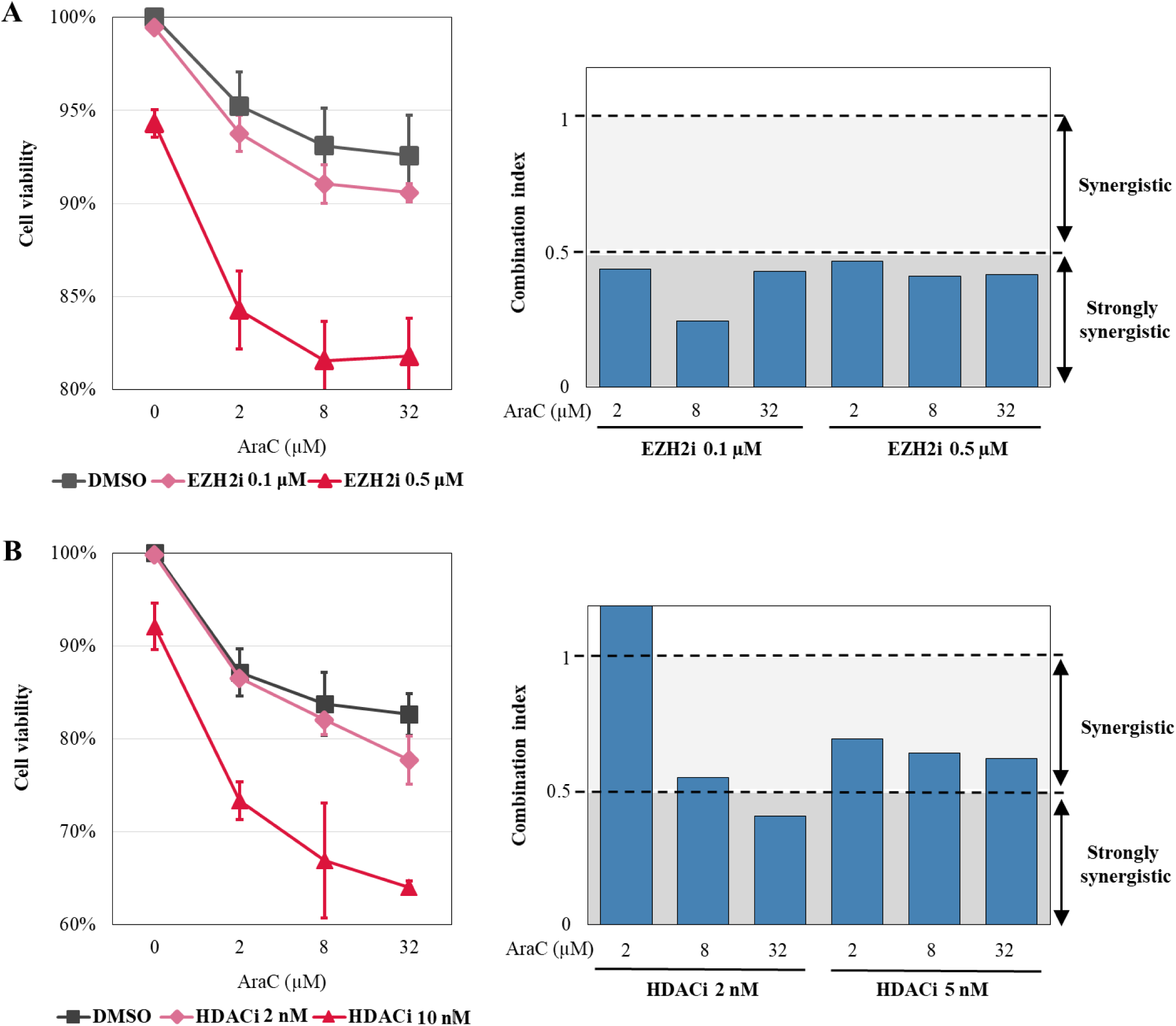
Combinatorial use of AraC with epigenetic modifiers has a synergistic effect. (A&B) Left: Cell viability evaluated by propidium iodide staining. SU-DHL6 cells were pretreated with tazemetostat (0.1µM or 0.5µM) for 4 days or panobinostat (2nM or 5nM) for 16 hours; then treated with the indicated concentrations of AraC (X-axis) for another 24 hours before cell viability was assessed by propidium iodide staining and Flow cytometry. Cell viability (%) was defined as the number of treated cells/untreated cells × 100 (Y-axis). Results are the average of three independent experiments with ± SD (standard deviation). Right: Combination index (CI) values assessed using the CompuSyn Software for data points of tazemetostat or panobinostat in combination with AraC. Shading represents the levels of synergism (> 1; synergy, > 0.5; strong synergy).

## DISCUSSION

In this study, we revealed that *SLFN11* expression is differentially regulated during B cell development. We found that SLFN11 is typically suppressed in GCBs (centroblasts and centrocytes) and GCB origin lymphomas. The suppression was achieved through histone methylation, and was reversible by the EZH2 inhibitor (tazemetostat) or the HDAC inhibitor (panobinostat). Pretreatment with these agents increased the cytocidal function of AraC and the combination could be applied to the treatment of GCB origin lymphomas.

Physiological reasons of why *SLFN11* needs to be suppressed in GCBs are not biologically examined in this study. However, in GCBs, activation-induced cytidine deaminase (AID) is specifically highly expressed to introduce somatic hypermutations in variable regions of immunoglobulin genes, whereas AID also induces DNA damages at non-target genes by generating apurinic sites [44]. Furthermore, GCBs proliferate very rapidly [42], which can enhance replication-dependent DNA damages [45]. Hence, GCBs are at high risk of DNA damage due to AID expression and rapid proliferation. Because SLFN11 exclusively kills replicating cells carrying genotoxic stress under the treatment of DNA damaging agents, we speculate that *SLFN11* needs to be downregulated in GCBs to avoid SLFN11-dependent cell death.

We then questioned which gene(s) suppresses SLFN11 at GCBs. We found that almost perfect inverse correlation between *SLFN11* expression and *PAX5*, a B-cell lineage-specific activator (Fig 1A and 1B). By mining a public database of chromatin immunoprecipitation-sequencing for PAX5 [46], we found a potential PAX5 binding site (GCGTGAC) in the promoter region of *SLFN11*, suggesting that PAX5 may be one of the repressors of *SLFN11* in B cells. This possibility is also supported by the fact that *SLFN11* expression is parallel to the expression of *PRDM1* and *XBP1*, which are the targets of PAX5 (Fig 1B).

Epigenetic regulation of *SLFN11* has been reported in other malignancies. In small cell lung cancer cells, *SLFN11* expression is silenced by marked deposition of H3K27me3, leading to drug resistance and recurrence after long treatment of platinum-derivatives, yet reactivated by inhibition of EZH2 a methyltransferase for H3K27 [25]. EZH2 inhibitor, tazemetostat has recently been approved by FDA for the treatment of follicular lymphoma [47]. Moreover, its efficacy for DLBCL is being studied [47]. In leukemia K562 and fibrosarcoma HT1080 cell lines, both of which have a very low basal SLFN11 expression, HDAC inhibitors (romidepsin and entinostat) increase SLFN11 expression and enhance sensitivity to DNA-damaging agents by SLFN11-dependent manner [26]. Our data consolidate these findings with GCB lymphoma lines and provide a rationale to treat B-cell lymphoma with low SLFN11 expression by the combination of tazometostat with AraC. Moreover, this is the first report showing that SLFN11 can be physiologically regulated through histone modifications during the normal developmental process.

As *SLFN11* is a promising target to sensitize tumor cells to cytotoxic chemotherapy, regulatory factors of SLFN11 expression are also favorable targets for the treatment [4, 48]. Our findings of dynamic regulation of SLFN11 during B-cell development will provide a basis to further investigate potential regulatory factors of SLFN11 at each developmental stage beyond PAX5 and histone modifiers.

## ACKNOWLEDGMENTS

We would like to thank A. Reddy and S. S. Dave, Duke University, for kindly providing access to the DLBCL database and F. Sasaki, Institute for Advanced Biosciences, Keio University, for technical assistance. This work was partly conducted through the Joint Research Program of the Radiation Biology Center, Kyoto University.

## Author contributions

Conceptualization: Fumiya Moribe, Momoko Nishikori, Junko Murai.

Methodology: Fumiya Moribe, Momoko Nishikori, Hiroyuki Sasanuma, Junko Murai.

Investigation: Fumiya Moribe, Remi Akagawa, Junko Murai.

Formal Analysis: Fumiya Moribe, Hiroshi Arima.

Writing – Original Draft Preparation: Fumiya Moribe, Momoko Nishikori, Junko Murai.

Supervision: Momoko Nishikori, Shunichi Takeda, Akifumi Takaori-Kondo, Junko Murai.

Funding Acquisition: Momoko Nishikori, Shunichi Takeda, Akifumi Takaori-Kondo, Junko Murai.

## Financial support

This work was supported by Grants-in-Aid for Scientific Research (JP18K08324 to M.N. and JP19H03505 to J.M.), a research grant from The Uehara Memorial Foundation (to J.M.), a Grant-in-Aid from the Ministry of Education, Science, Sport and Culture (KAKENHI 19K22561 and 16H06306 [to S.T.] and KAKENHI 18H04900 and 19H04267 [to H.S.]), the Japan Society for the Promotion of Science Core-to-Core Program, and Advanced Research Networks (to S.T.).

## Conflicts of interest

M.N. and A.T-K received honorarium and research funding from Eisai Co., Ltd. Other authors declare no conflicts of interest.

---

**Fig S1.**
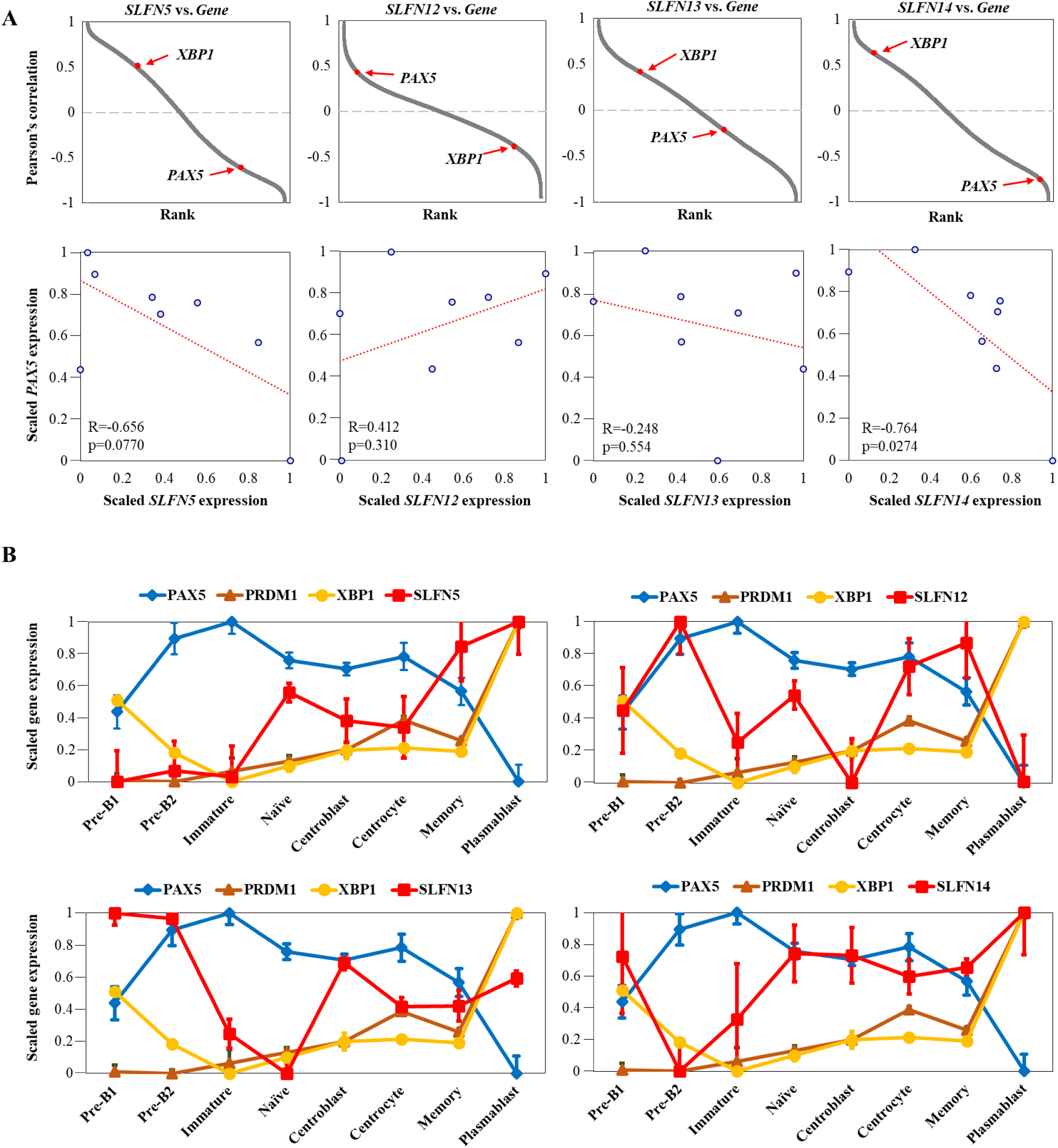
*SLFN* family members are differently expressed during B-cell development. (A) Upper: Pearson’s correlation between *SLFN* family members *(SLFN5, SLFN12, SLFN13, SLFN14)* and all the other genes. The genes are ordered from the highest correlation (left) to the lowest correlation (right). Lower: microarray gene expression plot of *SLFN* family members and *PAX5*. Precursor (Pre)-B1 cells, precursor (Pre)-B2 and immature B cells are were taken from human bone marrow (n=5), and naïve B cells, centroblasts, centrocytes, memory B cells and plasmablasts were taken from human tonsil (n=6). Pearson’s correlation (R), P-value (p) and an approximation straight line (a red dotted line) are shown. (B) Microarray gene expression profile (log2) of selected genes (*PAX5, PRDM1, XBP1, SLFN* family members) in human B cells from bone marrow and tonsil. Dots correspond to group means ± SE.

**Table S1. Primer sequences used for RT-qPCR**

**Table S2. Pearson’s correlation with *SLFN11* expression during B-cell development**

The data for the tables in this article are available from the link below.

https://data.mendeley.com/datasets/kn726mznn3/

